# Synergistic and Redundant Information Dynamics Exhibit Dissociable Alterations Across Major Psychiatric Disorders

**DOI:** 10.64898/2026.02.20.706948

**Authors:** Hinata Nago, Hiroki Kojima, Hiroyuki Yamaguchi, Yuichi Yamashita

## Abstract

Deficits in neural information integration are hypothesized to underlie diverse psychiatric symptoms, yet the specific patterns of alteration across different disorders remain unclear. In this study, we decomposed information dynamics between brain regions into synergistic and redundant components using a recent information-theoretical approach based on the idea of Partial Information Decomposition applied to resting-state fMRI data from patients with schizophrenia (SZ), autism spectrum disorder (ASD), and attention-deficit/hyperactivity disorder (ADHD). Our analysis revealed distinct disorder-specific profiles: SZ and ASD exhibited a widespread reduction in synergy, whereas ADHD showed a contrasting increase. Furthermore, ASD was uniquely characterized by a significant reduction in redundancy. Meta-analytic functional annotation using NeuroSynth associated synergy with higher-order cognitive functions and redundancy with lower-level sensorimotor processing. To investigate multivariate organization of these patterns that distinguish psychiatric diagnoses, we employed Linear Discriminant Analysis (LDA). This analysis demonstrated that synergy and redundancy partially capture distinct dimensions of network variation, exhibiting substantial complementarity in their multivariate structure. While redundancy overlapped considerably with correlation-based connectivity, synergy reflected additional structure not fully represented by conventional measures. Together, these findings indicate that decomposing information dynamics provides complementary perspectives on large-scale network organization, offering a refined framework for characterizing psychiatric disorders.

## 1 Introduction

The brain processes information from both the external world and internal states through a complex network of vast neural populations, giving rise to cognition and behavior. At the core of this sophisticated function lies ”information integration,” the process of consolidating inputs from multiple sources to generate integrated representations [1–4]. Neuroscience research has demonstrated that information integration—ranging from the binding of sensory modalities to global integration via fronto-posterior connections—is indispensable for establishing higher-order cognitive functions, such as memory, decision-making, and consciousness [5–8]. In parallel, psychiatric research suggests that deficits in information integration may underlie diverse psychiatric symptoms, such as the fragmentation of thought and perception in schizophrenia (SZ) or abnormalities in sensory integration in autism spectrum disorder (ASD) [9–11]. Although numerous neuroimaging studies have investigated information processing in the brain utilizing metrics such as functional connectivity (FC) [12] and transfer entropy [13, 14], these conventional methods fail to fully resolve the complex architecture of information integration. In addition, the quantitative assessment of information integration represents a critical challenge for advancing our understanding of both basic neuroscience and psychiatric pathophysiology.

In recent years, advanced analytical approaches have emerged to address this complexity. Artificial intelligence and machine learning techniques are increasingly applied to neuroimaging to capture latent disease-related representations [15, 16], and the field of ”Computational Psychiatry” has gained significant attention for its potential to bridge biological mechanisms and clinical symptoms [17–19]. However, despite their predictive power, these methods often fail to fully capture the information-theoretic interactions within brain networks, leaving the underlying structure of information dynamics—how information is processed and integrated across regions—largely unresolved.

To address this challenge, Partial Information Decomposition (PID) is gaining attention as a novel framework for quantitatively evaluating information integration [20]. PID decomposes the information provided by multiple sources to a target into interpretable components. Consider a scenario with two source variables and one target variable: information provided by only one source is defined as ”unique information”; information available from either source is ”redundant information (redundancy)”; and information cooperatively generated only when both sources are known is ”synergistic information (synergy)”. However, the information taxonomy introduced by classical PID is limited to scenarios with a single target variable, making it unable to capture how multiple variables collectively influence each other. This limitation prevents PID from providing an encompassing view of the dynamics of complex systems like the brain, where the past of multiple variables affects the future of multiple variables [2]. To address this, the framework has been extended to the time-series analysis of brain activity through Integrated Information Decomposition (ΦID) [2–4]. ΦID allows for the decomposition of information flow between brain regions into these redundant and synergistic components. This approach enables a comprehensive understanding of the qualitative aspects of information dynamics—distinguishing between robust, redundant pathways and complex, integrative synergistic processes—which conventional FC analyses, such as Pearson’s correlation, fail to capture [21].

In the context of psychiatric disorders, this decomposition offers new insights. For instance, a recent functional MRI (fMRI) study has suggested that in SZ, characterized by the fragmentation of thought and perception, ”synergy” within the brain is significantly reduced compared to healthy controls [22]. However, it remains unclear how ”synergy” and ”redundancy”—distinct constituent elements of information processing—are altered in other major psychiatric disorders, such as ASD and attention-deficit/hyperactivity disorder (ADHD), and how these alterations relate to specific cognitive domains.

Therefore, this study applies ΦID to resting-state fMRI data to comprehensively investigate brain-wide information dynamics across three major psychiatric disorders: SZ, ASD, and ADHD. Specifically, we identified patterns of alteration in ”synergy” and ”redundancy” in each patient group compared to healthy controls. Furthermore, our analysis extends beyond simple element-wise group comparisons by employing Linear Discriminant Analysis (LDA). Although univariate tests characterize distributed alterations across the brain, they do not explicitly describe how individual subjects are organized within a multivariate representational space. Given that psychiatric disorders are likely to reflect complex, large-scale network reconfigurations, examining their multivariate structure may provide complementary insights [23, 24]. Rather than focusing solely on classification performance, we employed LDA to explore how synergy, redundancy, and traditional correlation represent the relative relationships among diagnostic categories within a shared connectivity space. This approach allows us to examine whether decomposing information dynamics into synergistic and redundant components yields representational structures that differ from those obtained using standard correlation-based metrics.

Through these analyses, we aim to characterize how these information-theoretic metrics relate to distributed patterns of pathological variation across major psychiatric disorders. By contrasting these representations with correlation-based metrics, we assess whether synergy and redundancy provide complementary perspectives on brain organization. Rather than replacing standard metrics, this framework examines whether information decomposition can complement correlation-based connectivity analyses by capturing additional dimensions of network organization relevant to psychiatric evaluation.

## 2 Methods

### 2.1 Functional MRI datasets and preprocessing

Resting-state fMRI data were obtained from three publicly available databases: COBRE [https://fcon_1000.projects.nitrc.org/indi/retro/cobre.html] [25], ABIDE [https://fcon_1000.projects.nitrc.org/indi/abide/] [26], and ADHD-200 [https://fcon_1000.projects.nitrc.org/indi/adhd200/] [27]. The COBRE dataset included 71 patients with schizophrenia and 74 healthy controls. For the ABIDE and ADHD-200 datasets, we used participants acquired at the NYU site, consisting of 79 individuals with ASD and 105 typically developing controls in ABIDE, and 118 individuals with ADHD and 98 typically developing controls in ADHD-200.

Preprocessing was performed using the CONN toolbox ([https://www.nitrc.org/projects/conn/], version 22.v2407) following its standard default pipeline [28]. The steps included slice-timing correction, rigid-body realignment for head-motion correction, structural segmentation, and nonlinear spatial normalization to the MNI152 template using SPM12, followed by spatial smoothing with a 6-mm full-width at half-maximum (FWHM) Gaussian kernel [29]. Denoising was conducted using CONN’s default procedure, which incorporated regression of six motion parameters and their first derivatives, aCompCor components extracted from white-matter and cerebrospinal-fluid masks, and linear detrending [30]. BOLD time series were band-pass filtered between 0.008 and 0.09 Hz.

Following preprocessing, brain parcellation was performed using two atlases: (1) cortical regions were defined using the Schaefer-7-network functional atlas [31, 32], and (2) subcortical regions were defined using the Tian Subcortex Atlas (S2, 3T) [33]. These were combined to yield a total of 232 regions of interest (200 cortical and 32 subcortical), referred to as the Schaefer-232 brain atlas. Mean BOLD time series were extracted from each region and used for computing synergy and redundancy based on the ΦID framework [2–4].

### 2.2 Synergy and redundancy

#### 2.2.1 Partial Information Decomposition

We used the framework of Integrated Information Decomposition (ΦID) to calculate synergy and redundancy between two BOLD signal time series [2–4]. ΦID is grounded in Partial Information Decomposition (PID), which is an extension of Shannon’s information theory [20]. PID decomposes the mutual information between multiple sources and a single target into three types of information components: unique, redundant, and synergistic information.

Consider two source variables *X*_1_ and *X*_2_, and a target variable *Y* . According to the PID framework, the mutual information *I*(*X*_1_*, X*_2_; *Y*) can be decomposed as:

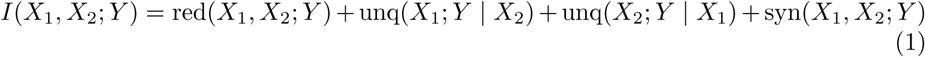

Here, red(*X*_1_*, X*_2_; *Y*) represents the information about *Y* that is shared by both *X*_1_ and *X*_2_. unq(*X*_1_; *Y* | *X*_2_) (and unq(*X*_2_; *Y* | *X*_1_)) denotes the information provided uniquely by *X*_1_ (or *X*_2_) about *Y* . Finally, syn(*X*_1_*, X*_2_; *Y*) corresponds to the information that is not provided by either *X*_1_ or *X*_2_ alone, but becomes accessible only when the source variables are observed jointly. Importantly, based on the inclusion-exclusion principle, defining a specific measure for redundancy allows the other information atoms (unique and synergistic information) to be automatically determined [20].

#### 2.2.2 Integrated Information Decomposition

However, the standard PID framework is restricted to single-target scenarios, making it unable to capture how multiple source variables collectively influence multiple target variables [2]. This limitation prevents PID from fully characterizing the dynamics of complex systems, such as the human brain, where the past states of multiple variables determine their future states. ΦID is a novel framework designed to overcome this limitation and is suitable for decomposing the time-delayed mutual information (TDMI) in a dynamic system [2–4].

Consider a system consisting of two time series ***X*** and ***Y*** (e.g., BOLD time series of two brain regions). Using the ΦID framework, the TDMI *I*(*X_t−τ_, Y_t−τ_* ; *X_t_, Y_t_*) can be decomposed into 16 ΦID atoms. This is because ΦID extends PID by simultaneously characterizing the information structure of both the sources (past) and targets (future), yielding 16 distinct modes of information flow (e.g., red→red, red→unq, etc.). In this study, we focused on ”synergy” corresponding to the atom ”syn→syn” and ”redundancy” corresponding to ”red→red” [2]. These quantities were computed using the open-source Python toolbox available at [https://github.com/Imperial-MIND-lab/integrated-info-decomp] [2–4]. Similar to PID, defining a specific measure for redundancy allows all other ΦID atoms to be derived. Specifically, we calculated the redundancy atom (red→red) using the Minimum Mutual Information (MMI) as follows [34]:

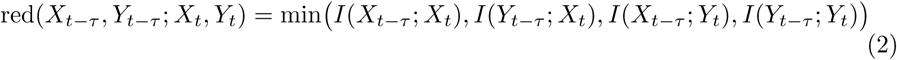

### 2.3 Traditional functional connectivity

Traditional functional connectivity was quantified using Pearson’s correlation between the BOLD signal time series of each pair of regions of interest (ROIs).

### 2.4 Statistical analysis of group differences

#### 2.4.1 Whole-brain analysis

We evaluated differences between patient and healthy control groups using independent two-sample *t*-tests for each metric: synergy, redundancy, and correlation. The *t*-statistics were defined such that a positive value (*t >* 0) indicates a reduction in the metric in patients compared to controls (i.e., Controls *>* Patients). To summarize regional abnormalities, the mean *t*-value for each ROI was calculated by averaging the *t*-values of all connections associated with that region.

#### 2.4.2 Network-level analysis

To evaluate differences in mean *t*-values across large-scale functional systems, ROIs were grouped into canonical functional networks. Specifically, we utilized the seven cortical networks defined by Yeo et al. (2011) [32] as implemented in the Schaefer atlas, supplemented by subcortical regions from the Tian atlas [31, 33]. The resulting networks include the visual (VIS), somatomotor (SOM), dorsal attention (DAN), salience/ventral attention (SAL), limbic (LIM), frontoparietal (FPN), and default mode (DMN) networks, as well as the subcortical network (SUB). Using these network assignments, we first performed Levene’s test to assess the homogeneity of variance among the network groups. We then conducted Welch’s one-way ANOVA to test for overall significant differences between networks, accounting for potential heteroscedasticity.

Following a significant main effect, post-hoc pairwise comparisons were performed between all pairs of networks using Welch’s *t*-tests. To control the family-wise error rate due to multiple testing, the resulting *p*-values were adjusted using the Holm-Bonferroni method.

### 2.5 NeuroSynth-based cognitive mapping analysis

While the canonical network-level analysis characterizes abnormalities in terms of intrinsic large-scale functional systems derived from resting-state organization, it does not directly specify the cognitive domains associated with the observed spatial patterns. To provide a complementary functional interpretation grounded in task-based evidence, we performed a NeuroSynth-based cognitive mapping of the *t*-value maps and synergy–redundancy gradients [https://neurosynth.org].

#### 2.5.1 Cognitive mapping of *t*-value–derived abnormality maps

For each disorder and metric, the *t*-values obtained from a two-sample *t*-test comparing patients and healthy controls for all brain region pairs were sorted in ascending order and divided into 5 percentile bins. ROI mask images were created from the group of brain region pairs within each bin, and these were used to identify the associated cognitive terms. We then calculated the spatial correlation between each mask and meta-analytic maps provided by NeuroSynth corresponding to 24 cognitive terms spanning lower-level sensorimotor to higher-order cognitive domains (e.g., visual perception, multisensory processing, social cognition), selected in accordance with previous studies [35, 36]. These correlations were converted into *z*-scores, and only terms with *z >* 3.1 were considered significantly associated with the identified brain regions. This analysis used a modified version of the original Python code published by Preti et al. (2019) [https://www.github.com/gpreti/GSP_StructuralDecouplingIndex] [36], adapted for the purposes of this study.

#### 2.5.2 Cognitive mapping of synergy–redundancy rank gradient

We also employed a ”synergy minus redundancy rank gradient” to define the ROI sets for NeuroSynth analysis, serving as a validation step to assess data integrity and reproduce hierarchical gradients reported in previous studies [3]. First, we calculated the nodal strength for each ROI by summing all connection weights in the group-averaged matrices for synergy and redundancy, respectively. ROIs were then ranked based on these nodal strength values in ascending order, such that the ROI with the lowest strength was assigned a rank of 1. The gradient was defined as the difference between the synergy rank and the redundancy rank for each ROI. All ROIs were sorted according to this gradient and divided into 5 percentile bins. Subsequent procedures, including the generation of ROI masks for each bin and the calculation of spatial correlations with NeuroSynth cognitive maps, were identical to those described in the *t*-value analysis section.

### 2.6 Linear Discriminant Analysis

#### 2.6.1 Diagnostic separability in multivariate connectivity space

While the preceding analyses focused on element-wise group differences and network-averaged summaries, they do not reveal how subjects are distributed within the high-dimensional representations derived from each information measure. To examine the multivariate organization of subjects in these representation spaces, we applied Linear Discriminant Analysis (LDA) separately to synergy, redundancy, and correlation. Input features were restricted to cortical regions; specifically, we extracted the upper-triangular elements of the connectivity matrix (excluding the diagonal), resulting in 19,900 unique connections. These features were then *z*-scored relative to the healthy control group.

Classification performance was evaluated using a 5-fold cross-validation scheme. To ensure a rigorous and unbiased comparison across the three metrics, the exact same partition of training and testing sets was applied to all modalities. In each fold, the LDA model was trained on the training set, and performance was quantified on the held-out test set. To provide a comprehensive evaluation of discriminability, we calculated three performance metrics: overall accuracy, balanced accuracy, and the macro-F1 score [37]. These values were averaged across the five folds to yield the final performance estimates.

Beyond classification accuracy, we examined the structure of the resulting low-dimensional space to understand how each metric separates diagnostic groups. The effect size of the separation was quantified as the difference between the centroids of the diagnostic groups along the canonical discriminant axes (LD1 and LD2). Specifically, we calculated the distance between the mean projected scores of the target groups to evaluate the clarity of the diagnostic structure encoded by each connectivity metric.

#### 2.6.2 Evaluation of residual site effects

To assess residual site effects in the *z*-scored features, we performed validation analyses using healthy controls from the COBRE, ABIDE, and ADHD-200 datasets [25–27]. Since the ABIDE and ADHD-200 samples were acquired at the same scanning site, they were combined into a single group. Consequently, site effects were evaluated by comparing two groups: the COBRE dataset versus the combined ABIDE/ADHD-200 dataset.

For each subject, connectivity matrices were summarized by computing the L2 norm of the upper-triangular elements. Differences between the two site groups were assessed using statistical comparisons, and effect sizes were quantified using Cohen’s *d*. This procedure was applied both before and after *z*-score normalization. In addition, the multivariate structure associated with potential site effects was examined using LDA. This analysis was applied to the healthy control data to visualize whether subjects clustered by imaging site after *z*-score normalization.

### 2.7 Analysis of discriminant weights

To interpret the specific connectivity patterns driving the diagnostic separation, we analyzed the discriminant weights (coefficients) assigned to the functional connections by the LDA model. The weight vectors corresponding to the canonical discriminant axes (LD1 and LD2) were extracted and reconstructed into 200 × 200 symmetric matrices, representing the contribution of each edge to the discrimination.

To summarize these patterns at a macro-scale, ROIs were grouped into 7 canonical functional networks [32]. We then quantified the contribution of each network pair (inter- and intra-network connections) using two metrics:

1. **Mean signed weight:** Calculated as the average of the raw weights within each network block. This metric indicates the directionality of the contribution (i.e., whether stronger connectivity pushes the projection score in the positive or negative direction).
2. **Mean absolute weight:** Calculated as the average of the absolute values of the weights within each network block. This metric quantifies the overall magnitude or importance of the network pair for the diagnostic discrimination, regardless of the direction of the effect.

### 2.8 Evaluation of prediction complementarity and oracle accuracy

To investigate whether the different information terms capture complementary diagnostic information, we analyzed the overlap of correct predictions between pairs of classifiers [38]. This analysis was performed on the pooled prediction labels obtained from the 5-fold cross-validation for all three pairwise combinations: synergy + redundancy, synergy + correlation, and redundancy + correlation.

For each pair of classifiers (denoted as model *A* and model *B*), we calculated the ”oracle accuracy”. This metric represents the theoretical maximum accuracy achievable by an ideal selector that correctly classifies a sample if at least one of the two models yields the correct prediction [38]. The oracle accuracy is defined as:

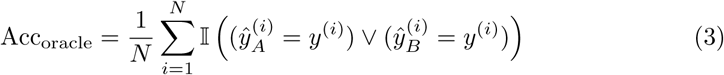

where *N* is the total number of samples, *y*^(*i*)^ is the true label for the *i*-th subject, 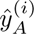 and 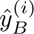 are the predicted labels by models *A* and *B*, respectively, and I(·) denotes the indicator function which equals 1 if the condition is true and 0 otherwise.

Furthermore, to quantify the statistical significance of the complementarity, we calculated the oracle gain, which measures the improvement over the best single model:

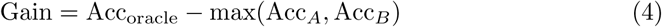

We estimated the 95% confidence intervals (CI) of the oracle gain using a bootstrapping procedure with 2,000 resamples. A gain was considered statistically significant if the lower bound of the 95% CI was greater than zero.

A higher oracle accuracy compared to the individual accuracy of each model suggests that the models err on different subsets of subjects. Consequently, the oracle gain serves as a quantitative indicator of complementarity, where a higher value signifies a greater improvement in classification performance achievable by combining the models. Additionally, to visualize the diversity of predictions, we computed the prediction disagreement rate and categorized the classification outcomes into four groups: (1) both correct, (2) only model *A* correct, (3) only model *B* correct, and (4) both wrong.

### 2.9 Correlation analysis with clinical severity

To investigate whether the decision boundaries identified by LDA captured clinically relevant variations within patient groups, we performed a post-hoc correlation analysis. For each information metric, we extracted the projected LDA scores (LD1 and LD2) for each subject.

We calculated Pearson correlation coefficients between the LDA scores and disorder-specific symptom severity ratings within each diagnostic group. Specifically, we analyzed 71 patients with SZ using the Positive and Negative Syndrome Scale (PANSS) [39], 79 patients with ASD using the Autism Diagnostic Observation Schedule (ADOS) [40], and 116 patients with ADHD using the Conners’ Parent Rating Scale-Revised: Long Version (CPRS-LV) [41]. Note that two subjects in the ADHD group were excluded from this specific analysis due to invalid symptom scores.

## 3 Results

### 3.1 Whole-Brain and network-level alterations in information dynamics

In this study, we applied the ΦID-based information decomposition approach to resting-state fMRI datasets [2–4] corresponding to three psychiatric disorders (SZ, ASD, and ADHD): COBRE (71 SZ patients, 74 healthy controls) [25], ABIDE (79 ASD patients, 105 healthy controls) [26], and ADHD-200 (118 ADHD patients, 98 healthy controls) [27]. Brain regions were defined based on the Schaefer-232 brain atlas (Fig. 1a) [31, 33]. For each participant, synergy and redundancy between the BOLD time series were calculated for all pairs of brain regions. Differences between patient and healthy control groups were assessed using independent *t*-tests. *t*-values were defined such that a positive value (*t >* 0) indicates a reduction in patients compared to controls. For comparison, *t*-values were similarly calculated for the traditional FC metric, Pearson’s correlation coefficient.

**Fig. 1:**
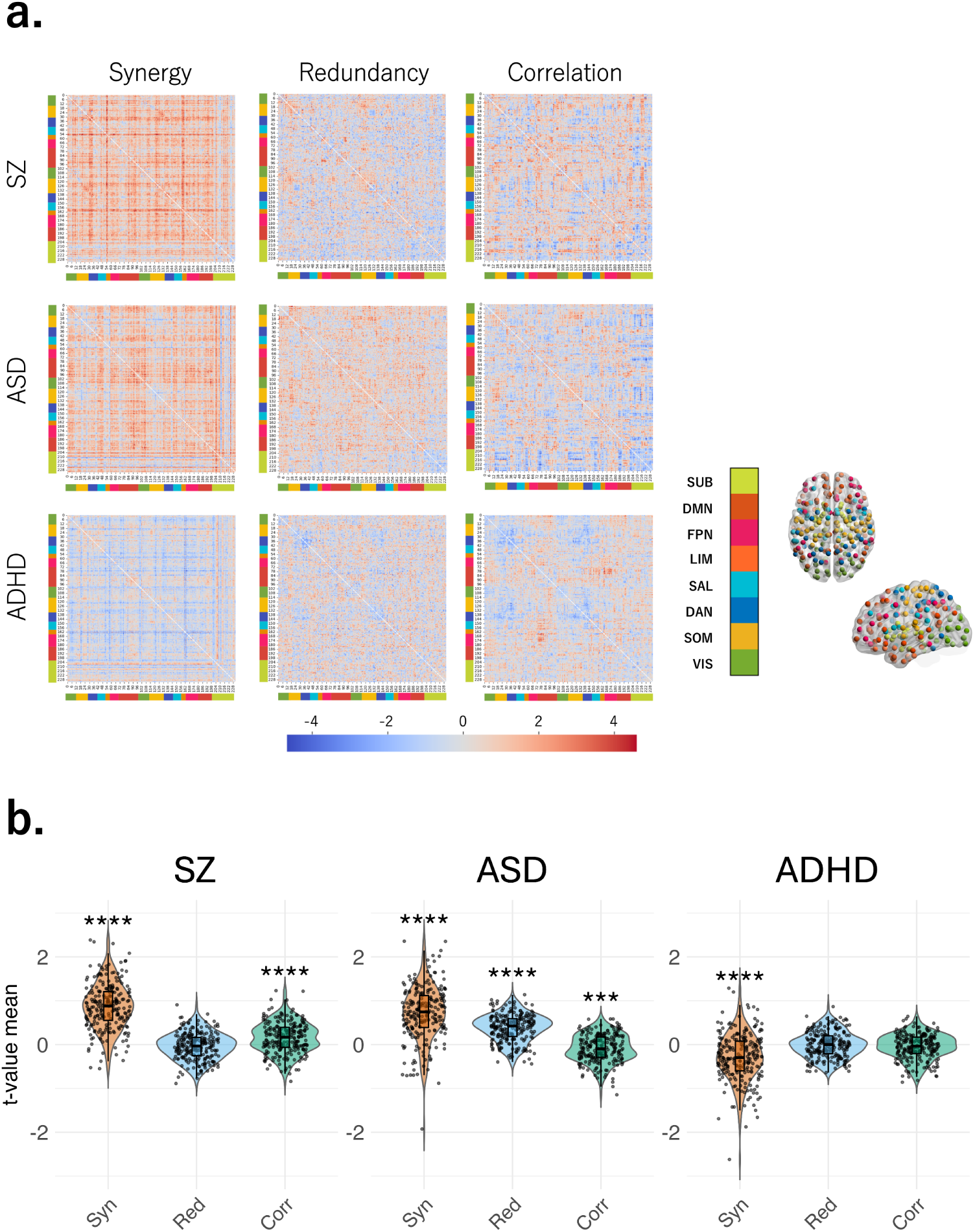
Whole-brain pattern comparison. **a.** Matrices of *t*-values derived from independent *t*-tests for each pair of brain regions, comparing patients with SZ, ASD, and ADHD to their respective healthy controls. Positive *t*-values indicate reduced connectivity in patients compared to controls. ROIs are organized by canonical functional networks [32]. Brain par-cellations were visualized using BrainNet Viewer [42]. **b.** The distribution of mean *t*-values across brain regions is shown for each metric (synergy, redundancy, correlation) and dis-order (SZ, ASD, ADHD). Internal box plots indicate the median and interquartile range, while individual data points represent the mean *t*-value of each ROI. Asterisks indicate a significant global deviation from zero (one-sample Wilcoxon signed-rank test, FDR corrected; \*\*\**p <* 0.001, \*\*\*\**p <* 0.0001).

We first visualized the *t*-value matrices of group differences to get an overview of the whole-brain alterations in information dynamics for each disorder (Fig. 1a). A characteristic pattern of widespread synergy reduction was observed in SZ, while in ASD, a reduction in both synergy and redundancy was noted. On the other hand, an increase in synergy was observed in the ADHD patient group. To quantify this overall trend, we plotted the distribution of mean *t*-values across brain regions for each metric (Fig. 1b). A widespread reduction in synergy was common to both SZ and ASD (*p <* 0.001), with ASD showing an additional prominent reduction in redundancy (*p <* 0.001). Conversely, in ADHD, an increase in synergy was observed in the patient group (*p <* 0.001). Notably, the reduction of synergy in SZ is consistent with the findings of a previous study [22]. Regarding traditional FC (correlation), significant reductions were observed in both SZ (*p <* 0.001) and ASD (*p <* 0.001).

To determine whether the whole-brain changes originated from specific functional networks, we next conducted a network-level analysis (Fig. 2). Comparing the mean *t*-values of each metric across 7 canonical functional networks based on the definition of Yeo et al. (2011) revealed distinct patterns for each disorder [32].

**Fig. 2:**
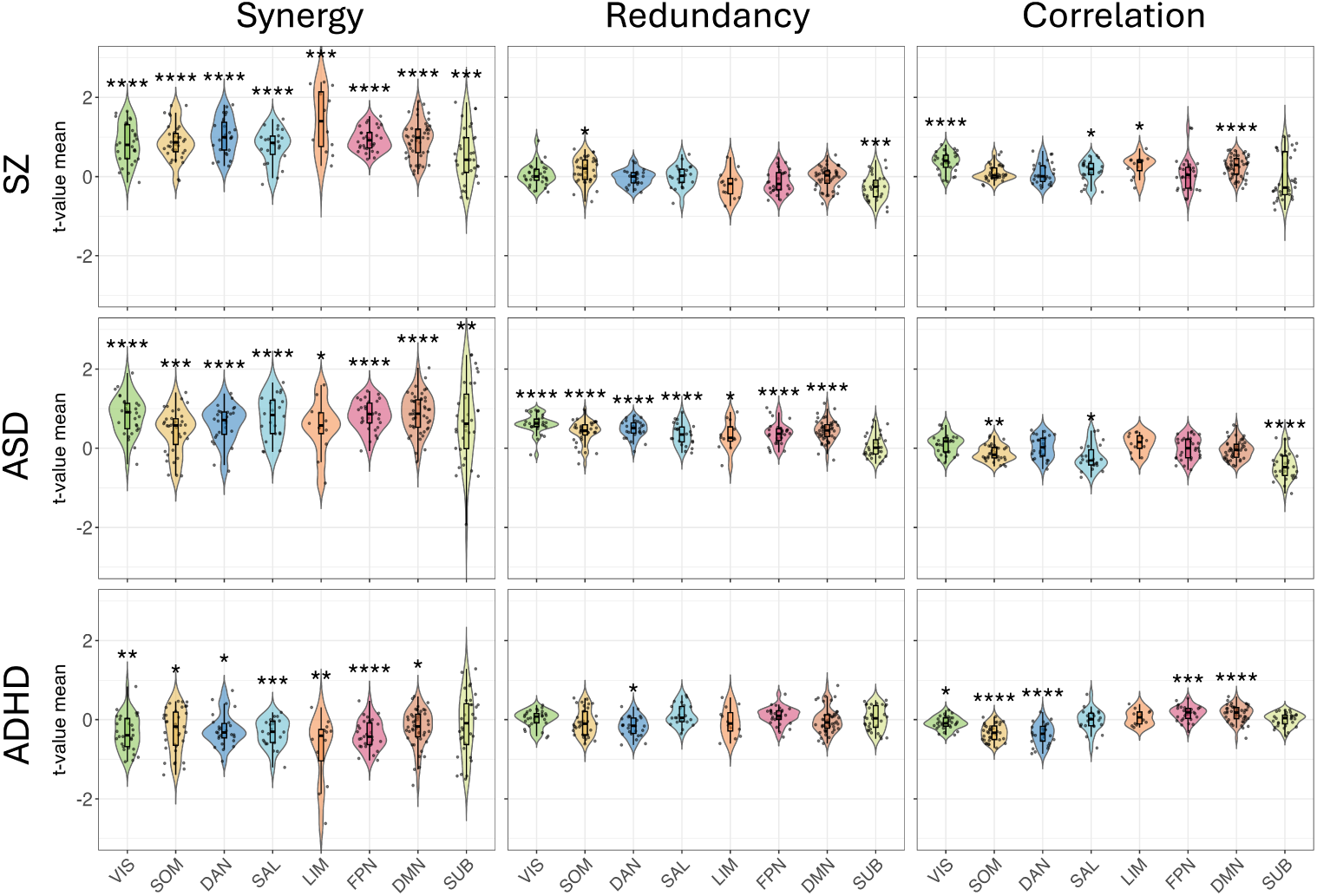
Network-level alterations in information decomposition metrics. Violin plots display the distributions of mean *t*-values stratified by canonical functional networks [32]. Rows represent the disorder groups (SZ, ASD, and ADHD), and columns represent the information metrics (synergy, redundancy, and correlation). Internal box plots indicate the median and interquartile range, while individual dots represent the mean *t*-values of ROIs belonging to each network. Asterisks indicate a significant global deviation from zero (one-sample Wilcoxon signed-rank test, FDR corrected; \**p <* 0.05, \*\**p <* 0.01, \*\*\**p <* 0.001, \*\*\*\**p <* 0.0001), indicating network-specific reductions (positive values) or increases (negative values) in patients compared to controls.

Specifically, in SZ, a significant reduction in synergy was observed across all networks (all *p <* 0.001). Furthermore, regions exhibiting the most notable reduction in synergy were frequently found in the limbic network (LIM). For this metric, ANOVA indicated significant differences in mean network values (Additional File, Table S1); however, subsequent post-hoc tests revealed no significantly different pairs (Additional File, Fig. S1). Regarding redundancy, significant alterations were limited to a decrease in the somatomotor network (SOM) (*p* = 0.024) and an increase in the subcortical network (SUB) (*p <* 0.001). Reduced correlation was observed in visual network (VIS), salience/ventral attention network (SAL), LIM, and default mode network (DMN) (all *p <* 0.05).

Similarly, synergy in ASD was reduced across all networks (all *p <* 0.05). ANOVA showed significant differences in network means, and post-hoc tests identified significant differences only in the SOM-frontoparietal network (FPN) and SOM-DMN pairs. Redundancy, in contrast, was reduced in all cortical networks (all *p <* 0.05) except for SUB. For redundancy, ANOVA indicated significant differences in network means (Additional File, Table S1), and post-hoc tests confirmed significant differences in the VIS-SOM, VIS-SAL, VIS-FPN, VIS-SUB, SOM-SUB, dorsal attention network (DAN)-SUB, SAL-SUB, FPN-SUB, and DMN-SUB pairs (Additional File, Fig. S1). Regarding correlation, there was a significant decrease in SOM and SAL (all *p <* 0.05), and an increase in SUB (*p <* 0.001).

In ADHD, synergy was increased, particularly within SAL and FPN networks (all *p <* 0.001). Regions showing a prominent increase were concentrated in LIM. However, ANOVA revealed no significant differences in network means (Additional File, Table S1), and no significant differences were found between any network pairs (Additional File, Fig. S1). For redundancy, only DAN exhibited significant alteration (*p* = 0.021). Lastly, there was a reduction in correlation in FPN and DMN (all *p <* 0.001), and increase in VIS, SOM, and DAN (all *p <* 0.05).

Furthermore, synergy exhibited greater variability across brain regions within the same network compared to the other two metrics (Additional File, Table S2).

Collectively, these results suggest that alterations in both information-theoretic metrics are not confined to specific canonical functional networks but rather exhibit widespread reductions (or increases) across the whole brain, although the degree of these alterations varies between the metrics.

### 3.2 Relationship between information dynamics changes and cognitive terms

While network-level analyses based on canonical functional networks situated the observed abnormalities within intrinsic large-scale functional systems derived from resting-state organization, this framework does not directly link these patterns to specific cognitive constructs established through task-based paradigms. To provide a complementary functional interpretation grounded in the task-based evidence, we conducted a NeuroSynth-based term-level analysis [35, 36, 43]. NeuroSynth is an automated meta-analytic tool that aggregates activation coordinates from thousands of published fMRI studies—including task-based paradigms—to statistically associate brain regions with cognitive terms. In this analysis, brain regions were stratified into percentiles based on their *t*-values, where higher percentiles correspond to larger *t*-values. We selected 24 topic terms based on previous studies, spanning the hierarchy from lower sensorimotor functions to higher-order cognitive domains [36, 43].

The results indicated that the information dynamics abnormalities observed in each disorder reflect impairments in distinct cognitive domains (Fig. 3). Consistent with the result of Fig. 1b, the transition point between positive and negative *t*-values deviated from the median in four metrics: synergy in SZ, synergy and redundancy in ASD, and synergy in ADHD. Brain regions with reduced synergy in SZ were strongly associated with cognitive terms such as visual semantics, visual perception, face/affective processing, and reward-based decision making. In ASD, regions with reduced synergy were closely related to reward-based decision making, inhibition, social cognition, and numerical cognition. On the other hand, regions with reduced redundancy in ASD were linked to sensory and perceptual processing domains, including visual perception, visuospatial processing, multisensory processing, and visual semantics. In ADHD, although the characteristic diagonal gradient from top-right to bottom-left was less distinct than in the other metrics, regions with increased synergy were closely related to terms such as visual perception, face/affective processing, emotion, and verbal semantics.

**Fig. 3:**
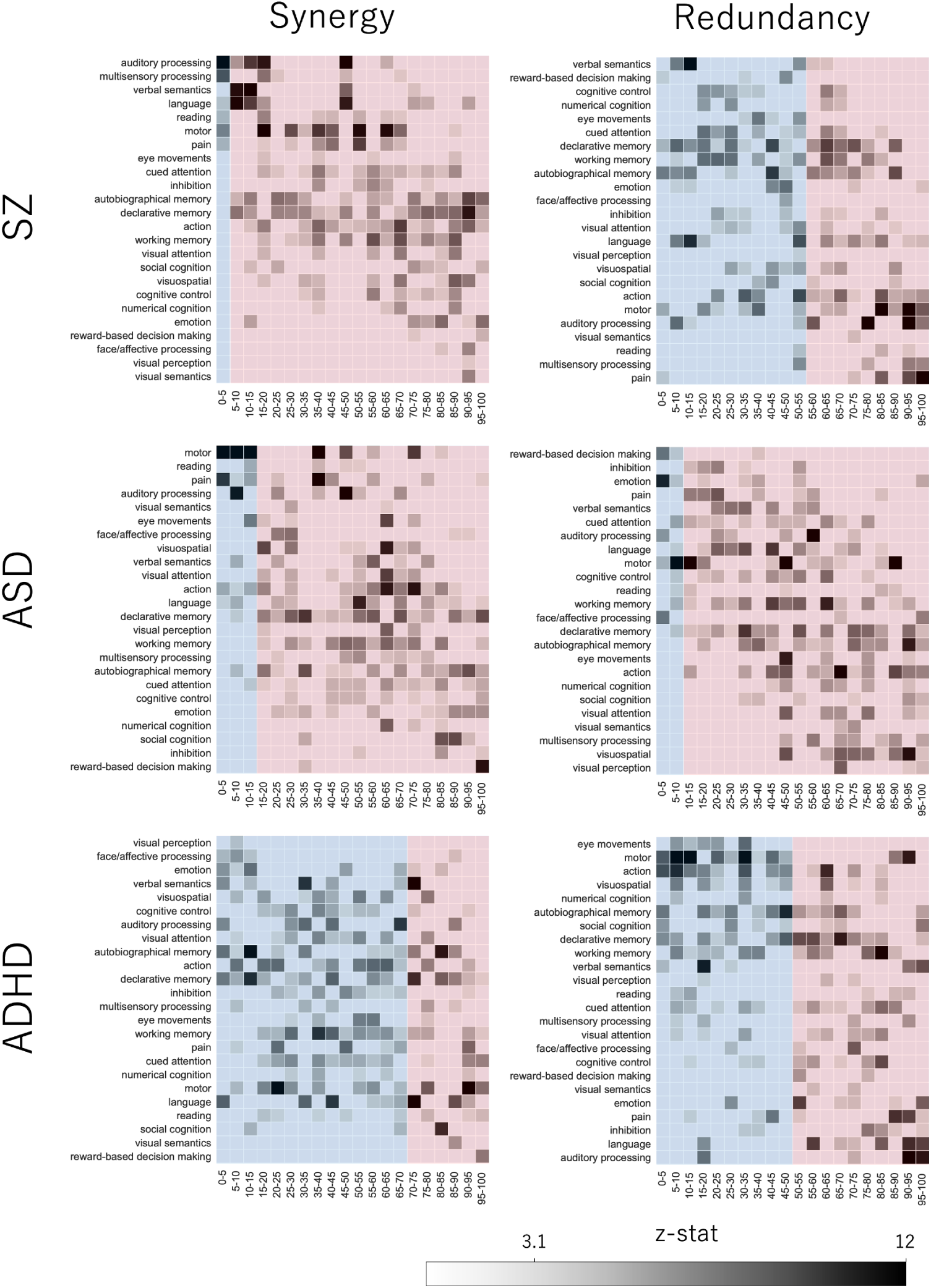
NeuroSynth-based analysis for each metric (synergy, redundancy) and disorder (SZ, ASD, ADHD). *t*-values for all brain region pairs were sorted in ascending order and divided into 5-percentile bins (X-axis). Higher percentiles correspond to larger *t*-values. The background colors denote the sign of the *t*-values: blue shading indicates negative values, while red shading indicates positive values. On the vertical axis, cognitive terms are arranged from top to bottom in ascending order of their centroid.

To ensure the reliability of our data, we also performed a validation analysis following previous studies using the synergy minus redundancy rank gradient (calculated by subtracting the redundancy rank from the synergy rank for each ROI) [3]. This analysis successfully reproduced the established trend where high-redundancy regions are associated with fundamental sensory processing, while high-synergy regions are linked to higher-order cognitive processes (Additional File, Fig. S2).

In summary, synergy dysfunction (both reduction and increase) is closely related to higher cognitive terms, while redundancy reduction is also strongly related to lower sensorimotor functions. These findings objectively demonstrate that the observed abnormalities in information dynamics for each disorder reflect impairments in different cognitive domains, providing a basis for the relationship with clinical symptoms discussed in the Discussion section.

### 3.3 LDA-based multivariate structure and clinical evaluation

We evaluated how each metric (synergy, redundancy, and correlation) captures the multivariate network organization characteristic of each psychiatric disorder. All features were converted into *z*-scores relative to healthy controls, ensuring that the resulting representations reflect normative deviations in network organization. Following the network-level analyses, subsequent multivariate analyses were restricted to cortical regions. To confirm that this normalization effectively mitigated potential scanner-related confounds, validation analyses were performed in healthy controls (comparing COBRE against ABIDE and ADHD-200, which originated from the same site). These analyses revealed significant site differences prior to normalization, which were effectively removed after *z*-score conversion; effect sizes were further reduced (all |Cohen’s *d* | *<* 0.03) and *p*-values increased (all *p >* 0.8), indicating negligible residual site effects across all three metrics (Additional File, Table S3). This assessment was further supported by LDA visualization, which revealed no site-specific clustering (Additional File, Fig. S3). To examine how subjects are organized within these multivariate representation spaces, we applied LDA separately to each connectivity modality (Fig. 4a). LDA was used as a supervised dimensionality-reduction approach that identifies axes maximizing between-group variance relative to within-group variance, thereby providing an interpretable low-dimensional summary of the relative configuration of diagnostic categories. In addition, we used the LDA projections of healthy controls as a qualitative check for residual site effects, confirming that subjects did not cluster by imaging site after *z*-score normalization (Additional File, Fig. S3). The overall classification accuracies for synergy, redundancy, and correlation were 0.608, 0.657, and 0.672, respectively. Comparable results were obtained using balanced accuracy and macro-F1 scores (Additional File, Table S4). Along the first linear discriminant axis (LD1), which primarily positioned SZ apart from the two neurodevelopmental disorders, synergy exhibited the largest effect size (2.13), followed by correlation (1.50) and redundancy (0.12). Similarly, on the second axis (LD2), which differentiated ASD from ADHD, synergy showed an effect size of 1.93, whereas correlation and redundancy recorded 1.41 and 0.19, respectively. These findings suggest that the dominant multivariate structure in the synergistic connectivity space most clearly reflected the relative configuration of diagnostic groups.

**Fig. 4:**
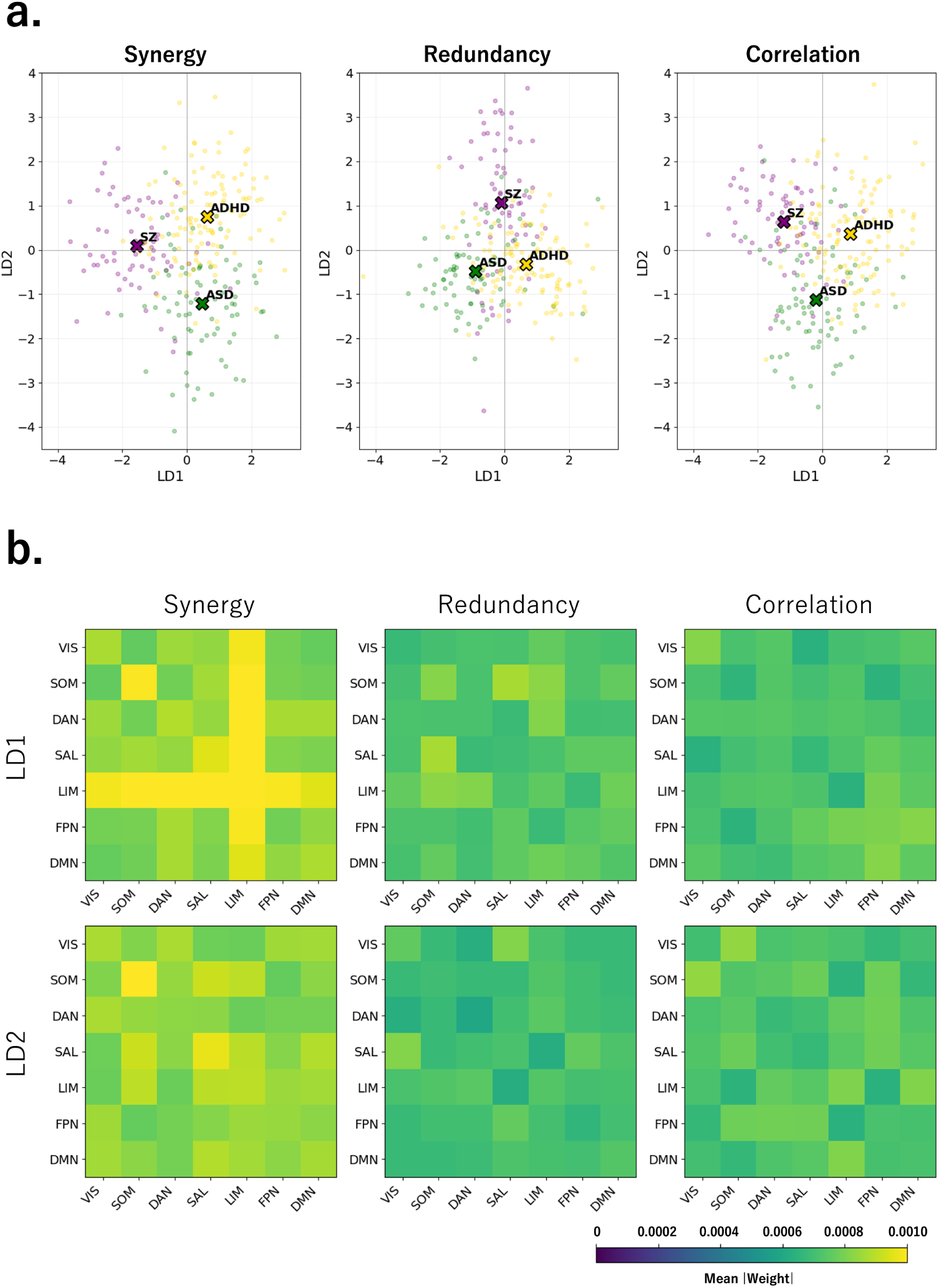
LDA of synergy, redundancy, and correlation in cortical regions. **a.** LDA projections of three diagnostic groups based on (left) synergy, (middle) redundancy, and (right) correlation derived from *z*-scored cortical data. Each dot represents an individual subject, color-coded by diagnosis (purple: SZ, green: ASD, yellow: ADHD). The ”×” markers indicate the centroids of each group. LD1 and LD2 represent the first and second discriminant axes, respectively. **b.** Displays the average of the absolute values of the LDA weights within each network block. This metric quantifies the overall magnitude or importance of each network pair for the diagnostic discrimination, regardless of the direction of the effect. The rows correspond to the first and second linear discriminants (LD1 and LD2).

To examine whether the LDA components of synergy, redundancy, and correlation captured individual variability in clinical severity, we correlated their LD1 and LD2 scores with disorder-specific symptom ratings: the Positive and Negative Syndrome Scale (PANSS) in SZ, the Autism Diagnostic Observation Schedule (ADOS) in ASD, and the Conners’ Parent Rating Scale-Revised: Long Version (CPRS-LV) in ADHD [39–41]. With the exception of a significant correlation between the LD2 score of redundancy and ADOS scores (*p* = 0.031), no significant associations were observed across the diagnostic groups (Additional File, Fig. S4). This indicates that the LDA dimensions primarily reflect between-group organization rather than within-group symptom variability, although redundancy features may partially relate to ASD severity.

### 3.4 Network-level structure of discriminant weights

To interpret the multivariate discriminant components in neurobiological terms, we projected the LDA weight matrices back to the ROI space and aggregated the absolute weights within and between functional networks, restricting the analysis to cortical regions. Differences in network-level structure were observed across the three modalities (Fig. 4b). While Fig. 2 depicts *t*-values reflecting patient–control differences, the LDA was performed on patient data *z*-scored relative to the healthy group. Accordingly, these discriminant weights reflect differences in multivariate organization between diagnostic groups rather than general deviations from normative connectivity. In the synergy-derived feature space, the LD1 weight map exhibited a pronounced concentration of contributions involving LIM. Inspection of the mean signed weights further indicated that these interactions were predominantly driven by negative coefficients (Additional File, Fig. S5 b). Because LD1 primarily positioned SZ relative to the other neurodevelopmental disorders, this pattern is consistent with the SZ-related synergy reductions concentrated in LIM observed in Fig. 2, where the most notable synergy reduction specific to SZ was concentrated in LIM. Intra-network connections within SOM also exhibited a relatively high absolute contribution. In contrast, both redundancy and correlation displayed more diffuse LD1 weight patterns, without a clear concentration in association networks. Although correlation showed moderate contributions within FPN and DMN, and redundancy within SOM and SAL, these patterns were less spatially differentiated than those observed for synergy.

A similar pattern was observed for LD2. Prominent LD2 weights were concentrated in SOM and SAL, and synergy generally exhibited higher mean absolute weights across connections compared to redundancy and correlation.

### 3.5 Complementarity of information-based representations

To investigate whether the distinct information-theoretic metrics capture complementary information for characterizing the three disorders (SZ, ASD, and ADHD), we analyzed the pairwise overlap of correct predictions between models. Table 1 summarizes the individual accuracies, oracle accuracies (theoretical maximums), and oracle gains (improvement over the best single model) for each pairwise combination [38]. Oracle accuracies for each diagnostic group are provided in Additional File, Table S5. The combination of synergy and redundancy demonstrated the most substantial diagnostic complementarity. While their individual classification accuracies were moderate (0.608 for synergy and 0.657 for redundancy), their combined oracle accuracy reached 0.828. This corresponds to a significant oracle gain of 0.170 (95% CI [0.127, 0.209]), the highest among all pairs. This result indicates that an ideal selector combining synergy and redundancy could improve predictive performance by 17% over the best single model. Furthermore, this pair exhibited a high prediction disagreement rate of 0.448, indicating that their misclassifications rarely overlapped. This suggests that synergy and redundancy capture partially non-overlapping patterns of normative deviation.

**Table 1:** Complementarity analysis and oracle accuracy across feature pairs.

| Pair (A + B) | Complementarity Metrics |  |  | Individual Accuracy |  |
| --- | --- | --- | --- | --- | --- |
|  | Oracle Gain [95% CI] | Oracle Acc. | Disagreement | Model A | Model B |
| **Syn + Red** | **0.170** [0.127, 0.209] | 0.828 | **0.448** | 0.608 | 0.657 |
| **Syn + Corr** | 0.163 [0.119, 0.205] | **0.836** | **0.448** | 0.608 | 0.672 |
| **Red + Corr** | 0.131 [0.097, 0.160] | 0.810 | 0.343 | 0.657 | 0.672 |

*Note.* Values represent accuracies and prediction disagreement rates. Oracle gain is shown with mean and 95% confidence intervals (CI).

A similar level of complementarity was observed for the synergy and correlation pair (oracle gain = 0.163, disagreement = 0.448), likely reflecting the fact that correlation partially encompasses synergistic information. In contrast, the redundancy and correlation pair showed the lowest complementarity (oracle gain = 0.131) and the lowest disagreement rate (0.343). This lower disagreement suggests that standard correlation and redundancy encode more overlapping aspects of network organization, whereas synergy provides a comparatively distinct contribution to the classification task. When all three metrics were combined, the oracle accuracy reached 0.892, with an oracle gain of 0.213.

Collectively, the high complementarity and prediction disagreement between synergy and redundancy indicate that these metrics capture partially distinct dimensions of network variation. These findings support the utility of decomposing correlation-based connectivity into information-theoretic components to reveal additional structure that is not fully reflected in conventional analyses.

## 4 Discussion

In this study, we employed a ΦID-based information decomposition approach to investigate patterns of information integration in the brain across three psychiatric disorders [2–4]. We revealed distinct alterations in information processing patterns specific to each disorder that were not fully reflected in traditional correlation-based connectivity measures. One of the key findings of this study is that impairments in information integration exhibit distinct patterns depending on the disorder. Specifically, SZ was characterized by a widespread reduction in synergy, whereas ASD showed reductions in both synergy and redundancy. In contrast, ADHD exhibited an increase in synergy. Notably, while traditional FC often highlights network-specific alterations, these information-theoretic metrics—particularly synergy—tended to show global, whole-brain shifts [23].

The widespread reduction in synergy observed in SZ replicates findings from a previous study and was prominent across functional networks [22]. From a complementary task-based perspective, NeuroSynth-based analysis indicated that this reduction in synergy is associated with higher-order cognitive functions such as visual semantics/perception and face/affective processing. Although synergy reductions were widespread across canonical networks, cognitive associations derived from the NeuroSynth database reflect an overlapping functional landscape in which individual regions may contribute to multiple cognitive domains. Accordingly, these functional interpretations are not intended as definitive one-to-one mappings, but rather as contextual indications of possible cognitive relevance. For completeness, we summarize the specific cognitive associations observed for each disorder below. In SZ, the observed reduction in synergy suggests that a disruption in the process of integrating fragmented information into a unified meaning may contribute to the diverse psychiatric symptoms observed in SZ, such as hallucinations, delusions, and negative symptoms [44]. In contrast, ASD revealed a unique pattern characterized by reductions in both synergy and redundancy. The reduction in synergy was associated with social cognition and inhibition, consistent with the deficits in social interaction that are core symptoms of ASD [45]. On the other hand, the reduction in redundancy was linked to fundamental sensory processing, such as visuospatial and multisensory processing. Since redundancy contributes to the robustness of the system against sensory inputs, this functional decline may relate to the neural basis of sensory hypersensitivity frequently reported in ASD [46]. Thus, ASD pathology may be conceptualized as a ”dual deficit” involving both impaired synergy (dysfunction of the social brain) and impaired redundancy (dysfunction of the sensory brain). In contrast, ADHD showed a whole-brain increase in synergy, excluding subcortical regions. Similar to SZ and ASD, regions with altered synergy in ADHD were preferentially associated with higher-order cognitive functions. One possible interpretation is that elevated synergy reflects altered modes of integrative processing that may support more flexible or divergent cognitive operations, which have been reported in individuals with ADHD [47, 48]. However, an alternative interpretation is that increased synergy reflects reduced network specialization or inefficient integration, rather than adaptive enhancement.

Although traditional correlation-based FC achieved higher classification accuracy, our LDA results suggest that information-theoretic metrics—particularly synergy—provide an alternative perspective on the multivariate organization of diagnostic groups. Specifically, synergy more clearly reflected the structural organization of psychopathology: LD1 positioned SZ relative to neurodevelopmental disorders (ASD/ADHD), while LD2 differentiated between ASD and ADHD. This pattern aligns with meta-analytic evidence from the ENIGMA consortium. Large-scale structural MRI studies have consistently reported that SZ exhibits widespread cortical thinning with larger effect sizes (Cohen’s *d*) compared to the more subtle and localized anatomical alterations observed in ASD and ADHD [49–52]. Our findings are consistent with the possibility that synergy reflects functional patterns corresponding to large-scale structural alterations observed in SZ.

The biological plausibility of this pattern is supported by the network-level distribution of discriminant weights. For LD1, which primarily positioned SZ relative to the other groups, synergy revealed a pronounced pattern with predominant weights concentrated in the LIM. LIM is central to emotion processing, motivation, and salience attribution—functions heavily impaired in SZ [53]. The concentration of synergy reduction in the LIM is consistent with the ”aberrant salience” hypothesis and previous findings of limbic dysconnectivity [9, 54–56]. Furthermore, the LD2 axis provided insights into the relative positioning of ASD and ADHD. In the synergy-based representation, this relationship was associated with weights in SOM and SAL. This result is particularly meaningful when considering the ”local vs. global” contrast: ADHD is characterized by widespread, diffuse increases in synergy, whereas ASD exhibits more focal deviations [57, 58]. The high contributions of SOM and SAL may reflect ASD-related alterations in sensory-motor integration and salience processing, allowing the model to distinguish ASD from the globally altered reference of ADHD [59, 60].

The complementarity analysis provided quantitative support for the functional dissociation discussed above. The combination of synergy and redundancy yielded the highest oracle gain, indicating that these two metrics capture partially non-overlapping aspects of network variation. This statistical independence aligns with the distinct information-theoretic profiles of the disorders. Specifically, while both SZ and ASD are characterized by a global reduction in synergy (in contrast to the global increase observed in ADHD), ASD is uniquely distinguished by a widespread reduction in redundancy. Consequently, this ASD-specific deficit in redundancy may have contributed to the differentiation between SZ and ASD. By combining these orthogonal signals, the model provides a ”dual-view” of network organization. In contrast, standard correlation showed low complementarity with redundancy, suggesting that traditional FC largely encodes redundant interactions and fails to explicitly capture the synergistic components of information processing. This interpretation is consistent with prior theoretical work demonstrating that redundancy substantially contributes to classic statistical dependencies, including covariance, precision, and mutual information [21]. This observation highlights a limitation of conventional analyses: by aggregating synergistic and redundant dynamics into a single correlation metric, distinct network patterns may be conflated. Therefore, our findings support the value of decomposing correlation-based FC into its information-theoretic constituents to reveal additional dimensions of network organization.

Our study has several limitations. First, potential confounding effects arising from the multi-site nature of the datasets cannot be fully ruled out. Second, it remains unclear how the qualitative difference between increased and decreased synergy maps onto specific clinical abnormalities. Moreover, establishing robust associations between these information-theoretic metrics and objective symptom scores or quantitative behavioral phenotypes remains a critical goal for future investigation. Finally, bridging macroscopic information dynamics with microscopic neural mechanisms remains a challenge for future research.

Despite these limitations, this study demonstrates that decomposing information dynamics into synergistic and redundant components reveals distinct patterns that are not fully captured by correlation analysis. We identified that schizophrenia is characterized by a global reduction of synergistic integration, whereas ASD presents a ”dual deficit” in both social (synergy) and sensory (redundancy) processing. ADHD exhibits a brain-wide increase in synergy. By disentangling these complementary signals, our findings highlight the utility of information decomposition as a refined framework for characterizing network-level alterations in psychiatric disorders.

## Abbreviations

ADHD: Attention-deficit/hyperactivity disorder
ADOS: Autism Diagnostic Observation Schedule
ASD: Autism spectrum disorder
CI: Confidence intervals
Corr: Correlation
CPRS-LV: Conners’
Parent Rating Scale-Revised: Long Version
DAN: Dorsal attention network
DMN: Default mode network
FC: Functional connectivity
fMRI: Functional magnetic resonance imaging
FPN: Frontoparietal network
LDA: Linear Discriminant Analysis
LIM: Limbic network
PANSS: Positive and Negative Syndrome Scale
PID: Partial Information Decomposition
Red: Redundancy
ROI: Region of interest
SAL: Salience/ventral attention network
SOM: Somatomotor network
SUB: Subcortical network
Syn: Synergy
SZ: Schizophrenia
TDMI: Time-delayed mutual information
VIS: Visual network
ΦID: Integrated Information Decomposition

## 5 Declarations

### 5.1 Availability of data and materials

The resting-state fMRI datasets analyzed in this study are publicly available from the following repositories: COBRE [https://fcon_1000.projects.nitrc.org/indi/retro/cobre.html] [25], ABIDE [https://fcon_1000.projects.nitrc.org/indi/abide/] [26], and ADHD-200 [https://fcon_1000.projects.nitrc.org/indi/adhd200/] [27]. For data preprocessing and analysis, the CONN toolbox is available at https://www.nitrc.org/projects/conn/ [28]. The Python code used to calculate synergy and redundancy is openly available from https://github.com/Imperial-MIND-lab/integrated-info-decomp [2, 3]. Additionally, the Python scripts used for the NeuroSynth-based analysis can be found at https://github.com/gpreti/GSP_StructuralDecouplingIndex [36].

### 5.2 Competing interests

The authors declare that they have no competing interests.

### 5.3 Funding

This study was partly supported by AMED Multidisciplinary Frontier Brain and Neuroscience Discoveries (Brain/MINDS 2.0) JP24wm0625407, JST CREST JPMJCR21P4, JSPS KAKENHI JP22K15777, JP24H00076, JP24K00499, JP25H01173, and Intramural Research Grant (6-9, 7-9) for Neurological and Psychiatric Disorders of NCNP.

### 5.4 Authors’ contributions

H.N., H.K., and Y.Y. conceived the original idea for the study. H.Y. performed the fMRI data preprocessing. H.N. analyzed the data and drafted the manuscript. All authors provided critical feedback and approved the final manuscript.

## Acknowledgements

Not applicable.

## Notes

https://github.com/Imperial-MIND-lab/integrated-info-decomp

### Competing Interest Statement

The authors have declared no competing interest.

