## Additional file for "Synergistic and Redundant Information Dynamics Exhibit Dissociable Alterations Across Major Psychiatric Disorders"

**Table S1**: Welch's One-Way ANOVA across canonical functional networks.

| | **Synergy** | | **Redundancy** | | **Correlation** | |
|---|---|---|---|---|---|---|
| | ***F*-value** | ***p*-value** | ***F*-value** | ***p*-value** | ***F*-value** | ***p*-value** |
| **SZ** | 2.64 | 0.017 | 6.79 | <0.001 | 5.56 | <0.001 |
| **ASD** | 2.71 | 0.014 | 10.62 | <0.001 | 10.17 | <0.001 |
| **ADHD** | 1.13 | 0.351 | 3.17 | 0.005 | 27.41 | <0.001 |

**Table S2**: Levene's Test for homogeneity of variances across canonical functional networks.

| | **Synergy** | | **Redundancy** | | **Correlation** | |
|---|---|---|---|---|---|---|
| | ***F*-value** | ***p*-value** | ***F*-value** | ***p*-value** | ***F*-value** | ***p*-value** |
| **SZ** | 4.56 | <0.001 | 1.22 | 0.294 | 5.15 | <0.001 |
| **ASD** | 4.85 | <0.001 | 0.69 | 0.679 | 1.62 | 0.132 |
| **ADHD** | 4.76 | <0.001 | 1.83 | 0.082 | 1.94 | 0.065 |

**SZ — Synergy**

| | VIS | SOM | DAN | SAL | LIM | FPN | DMN | SUB |
|---|---|---|---|---|---|---|---|---|
| VIS | | | | | | | | |
| SOM | 1 | | | | | | | |
| DAN | 1 | 1 | | | | | | |
| SAL | 1 | 1 | 1 | | | | | |
| LIM | 0.673 | 0.837 | 1 | 0.451 | | | | |
| FPN | 1 | 1 | 1 | 1 | 1 | | | |
| DMN | 1 | 1 | 1 | 1 | 1 | 1 | | |
| SUB | 1 | 0.521 | 0.056 | 1 | 0.084 | 0.102 | 0.172 | |

**SZ — Redundancy**

| | VIS | SOM | DAN | SAL | LIM | FPN | DMN | SUB |
|---|---|---|---|---|---|---|---|---|
| VIS | | | | | | | | |
| SOM | 1 | | | | | | | |
| DAN | 1 | 0.093 | | | | | | |
| SAL | 1 | 0.895 | 1 | | | | | |
| LIM | 0.895 | 0.115 | 1 | 1 | | | | |
| FPN | 0.704 | 0.012 | 1 | 1 | 1 | | | |
| DMN | 1 | 0.122 | 1 | 1 | 1 | 1 | | |
| SUB | 0.003 | <0.001 | 0.024 | 0.032 | 1 | 0.895 | 0.017 | |

**SZ — Correlation**

| | VIS | SOM | DAN | SAL | LIM | FPN | DMN | SUB |
|---|---|---|---|---|---|---|---|---|
| VIS | | | | | | | | |
| SOM | <0.001 | | | | | | | |
| DAN | 0.004 | 1 | | | | | | |
| SAL | 0.081 | 1 | 1 | | | | | |
| LIM | 1 | 0.931 | 1 | 1 | | | | |
| FPN | 0.033 | 1 | 1 | 1 | 1 | | | |
| DMN | 0.859 | 0.033 | 0.280 | 1 | 1 | 0.714 | | |
| SUB | 0.018 | 1 | 1 | 1 | 0.530 | 1 | 0.267 | |

**ASD — Synergy**

| | VIS | SOM | DAN | SAL | LIM | FPN | DMN | SUB |
|---|---|---|---|---|---|---|---|---|
| VIS | | | | | | | | |
| SOM | 0.093 | | | | | | | |
| DAN | 1 | 1 | | | | | | |
| SAL | 1 | 0.551 | 1 | | | | | |
| LIM | 1 | 1 | 1 | 1 | | | | |
| FPN | 1 | 0.035 | 1 | 1 | 1 | | | |
| DMN | 1 | 0.012 | 0.748 | 1 | 1 | 1 | | |
| SUB | 1 | 1 | 1 | 1 | 1 | 1 | 1 | |

**ASD — Redundancy**

| | VIS | SOM | DAN | SAL | LIM | FPN | DMN | SUB |
|---|---|---|---|---|---|---|---|---|
| VIS | | | | | | | | |
| SOM | 0.050 | | | | | | | |
| DAN | 0.524 | 1 | | | | | | |
| SAL | 0.020 | 1 | 1 | | | | | |
| LIM | 0.387 | 1 | 1 | 1 | | | | |
| FPN | 0.012 | 1 | 1 | 1 | 1 | | | |
| DMN | 0.107 | 1 | 1 | 1 | 1 | 1 | | |
| SUB | <0.001 | <0.001 | <0.001 | 0.010 | 0.634 | 0.001 | <0.001 | |

**ASD — Correlation**

| | VIS | SOM | DAN | SAL | LIM | FPN | DMN | SUB |
|---|---|---|---|---|---|---|---|---|
| VIS | | | | | | | | |
| SOM | 0.005 | | | | | | | |
| DAN | 1 | 0.369 | | | | | | |
| SAL | 0.004 | 1 | 0.089 | | | | | |
| LIM | 1 | 0.094 | 1 | 0.018 | | | | |
| FPN | 1 | 0.555 | 1 | 0.139 | 1 | | | |
| DMN | 0.237 | 0.961 | 1 | 0.241 | 0.555 | 1 | | |
| SUB | <0.001 | 0.001 | <0.001 | 0.369 | <0.001 | <0.001 | <0.001 | |

**ADHD — Synergy**

| | VIS | SOM | DAN | SAL | LIM | FPN | DMN | SUB |
|---|---|---|---|---|---|---|---|---|
| VIS | | | | | | | | |
| SOM | 1 | | | | | | | |
| DAN | 1 | 1 | | | | | | |
| SAL | 1 | 1 | 1 | | | | | |
| LIM | 1 | 1 | 1 | 1 | | | | |
| FPN | 1 | 1 | 1 | 1 | 1 | | | |
| DMN | 1 | 1 | 1 | 1 | 1 | 1 | | |
| SUB | 1 | 1 | 1 | 1 | 1 | 1 | 1 | |

**ADHD — Redundancy**

| | VIS | SOM | DAN | SAL | LIM | FPN | DMN | SUB |
|---|---|---|---|---|---|---|---|---|
| VIS | | | | | | | | |
| SOM | 1 | | | | | | | |
| DAN | 0.121 | 1 | | | | | | |
| SAL | 1 | 0.532 | 0.010 | | | | | |
| LIM | 1 | 1 | 1 | 1 | | | | |
| FPN | 1 | 0.682 | 0.006 | 1 | 1 | | | |
| DMN | 1 | 1 | 0.719 | 1 | 1 | 1 | | |
| SUB | 1 | 1 | 0.218 | 1 | 1 | 1 | 1 | |

**ADHD — Correlation**

| | VIS | SOM | DAN | SAL | LIM | FPN | DMN | SUB |
|---|---|---|---|---|---|---|---|---|
| VIS | | | | | | | | |
| SOM | <0.001 | | | | | | | |
| DAN | 0.001 | 1 | | | | | | |
| SAL | 1 | 0.001 | 0.001 | | | | | |
| LIM | 0.417 | <0.001 | <0.001 | 1 | | | | |
| FPN | <0.001 | <0.001 | <0.001 | 0.208 | 1 | | | |
| DMN | <0.001 | <0.001 | <0.001 | 0.208 | 1 | 1 | | |
| SUB | 0.531 | <0.001 | <0.001 | 1 | 1 | 0.013 | 0.010 | |

**Fig. S1**: Post-hoc pairwise statistical comparisons across canonical functional networks. Each table displays the $p$-values resulting from post-hoc pairwise comparisons using Welch's t-tests for synergy, redundancy, and correlation across the three disorder groups (SZ, ASD, and ADHD) . The resulting $p$-values were adjusted using the Holm-Bonferroni method. Cells highlighted in orange indicate statistically significant differences ($p < 0.05$).

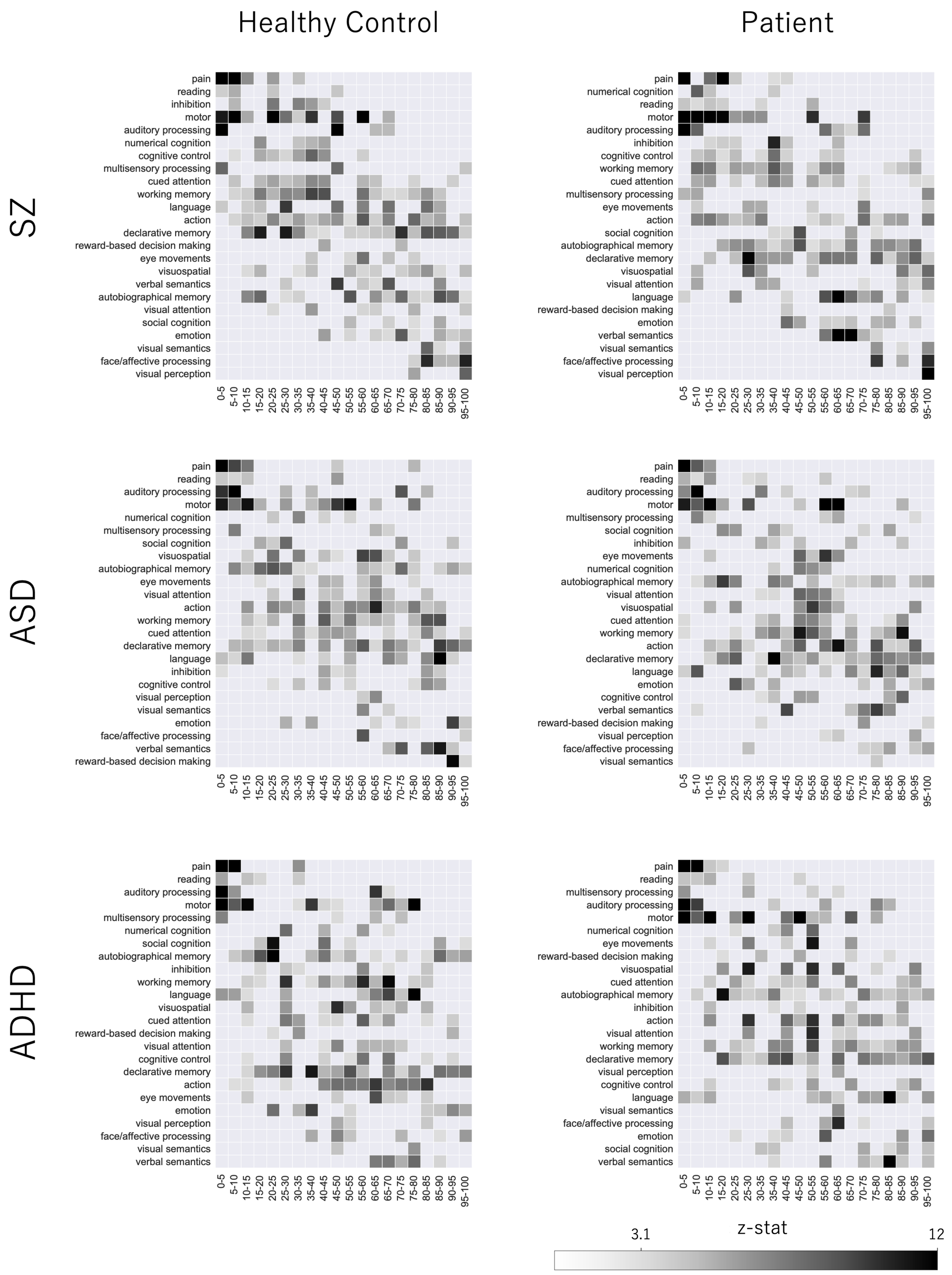


**Fig. S2**: NeuroSynth term-based meta-analysis using the synergy-redundancy rank gradient. Meta-analytic mapping illustrates the spatial association between brain regions and cognitive terms, ordered by the synergy minus redundancy rank gradient. The analysis was performed separately for healthy controls and patient groups (SZ, ASD, and ADHD) across 5-percentile bins .

**Table S3**: Statistical assessment of residual site effects in healthy controls before and after z-score normalization.

| | **Synergy** | | **Redundancy** | | **Correlation** | |
|---|---|---|---|---|---|---|
| | ***p*-value** | **Cohen's *d*** | ***p*-value** | **Cohen's *d*** | ***p*-value** | **Cohen's *d*** |
| **Raw** | 0.892 | -0.02 | 0.238 | 0.16 | 0.269 | 0.15 |
| ***Z*-scored** | 0.964 | -0.01 | 0.950 | 0.02 | 0.864 | 0.02 |

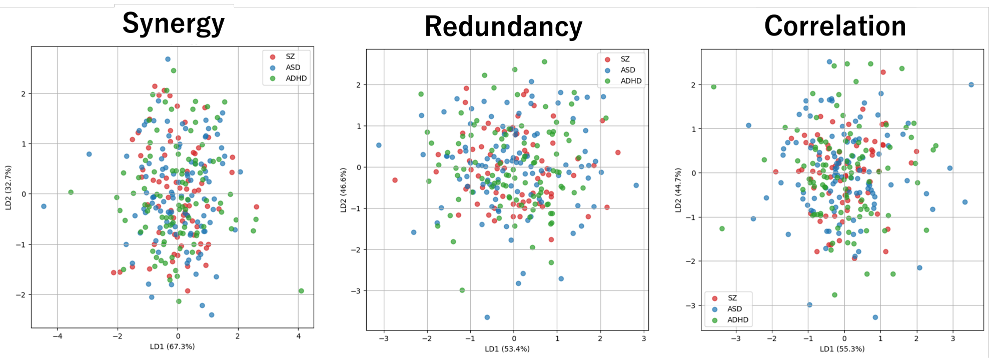


**Fig. S3**: LDA for $z$-score normalized healthy controls. Scatter plots visualize the first two discriminant axes (LD1 and LD2) for synergy, redundancy, and correlation using z-scored connectivity features of healthy controls.

**Table S4**: Classification performance of LDA models across connectivity modalities.

| **Metric** | **Synergy** | **Redundancy** | **Correlation** |
|---|---|---|---|
| Overall Accuracy | 0.608 | 0.657 | 0.672 |
| Balanced Accuracy | 0.581 | 0.601 | 0.652 |
| Macro-F1 Score | 0.589 | 0.608 | 0.659 |

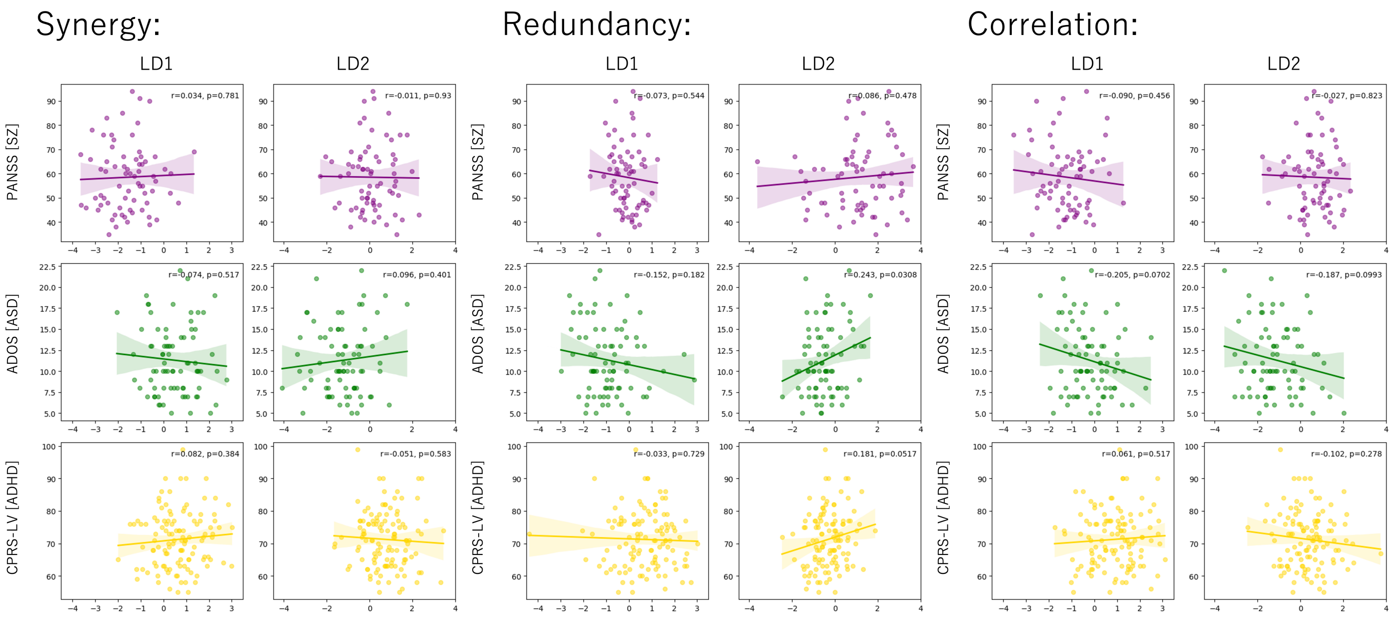


**Fig. S4**: Correlation between LDA projected scores and clinical symptom severity. Scatter plots display the relationships between individual LD1 and LD2 scores for each connectivity metric (synergy, redundancy, correlation) and disorder-specific clinical symptom scales (PANSS for SZ, ADOS for ASD, and CPRS-LV for ADHD) .

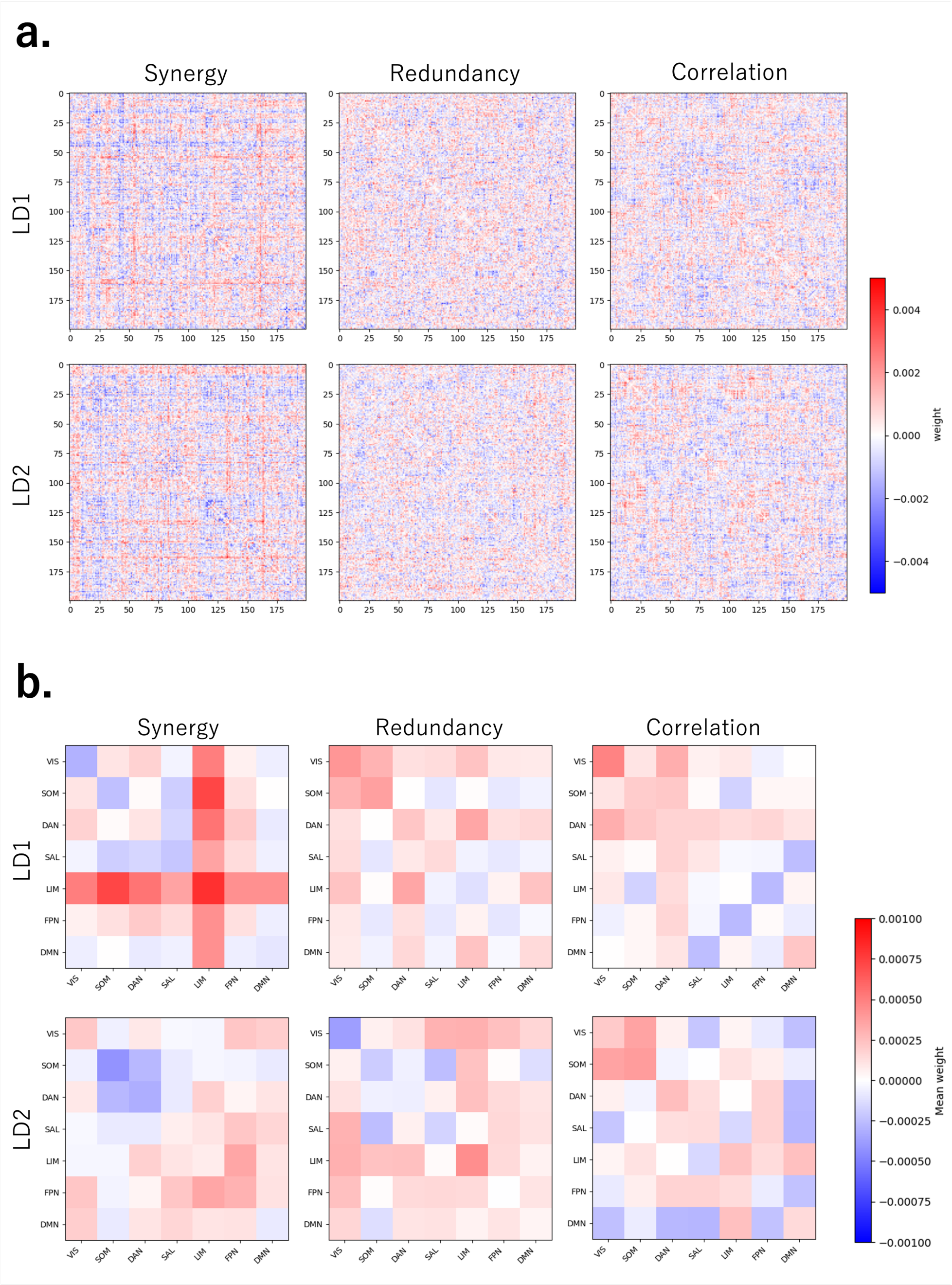


**Fig. S5**: Distribution of LDA weights. **a.** Reconstructed $200 \times 200$ symmetric matrices representing the raw discriminant weights for LD1 and LD2 . **b.** Mean signed weights aggregated within and between canonical functional networks .

**Table S5**: Group-wise classification performance and non-redundant contribution counts across feature pairs.

| Pair (A + B) | Group | $n$ | A Acc. | B Acc. | Oracle Acc. | Only A | Only B |
|---|---|---|---|---|---|---|---|
| **Syn + Red** | SZ | 71 | 0.507 | 0.296 | 0.634 | 24 | 9 |
| | ASD | 79 | 0.481 | 0.582 | 0.785 | 16 | 24 |
| | ADHD | 118 | 0.754 | 0.924 | 0.975 | 6 | 26 |
| **Syn + Corr** | SZ | 71 | 0.507 | 0.563 | 0.746 | 13 | 17 |
| | ASD | 79 | 0.481 | 0.620 | 0.797 | 14 | 25 |
| | ADHD | 118 | 0.754 | 0.771 | 0.915 | 17 | 19 |
| **Red + Corr** | SZ | 71 | 0.296 | 0.563 | 0.662 | 7 | 26 |
| | ASD | 79 | 0.582 | 0.620 | 0.722 | 8 | 11 |
| | ADHD | 118 | 0.924 | 0.771 | 0.958 | 22 | 4 |

*Note.* "Only A" and "Only B" columns represent the number of subjects correctly classified by only that specific metric.
